# ChiMER: Integrating chromatin architecture into splicing graphs for chimeric enhancer RNAs detection

**DOI:** 10.64898/2026.03.16.711958

**Authors:** Yujia Xiang, Xinyu Xiao, Bingkun Zhou, Linhai Xie

## Abstract

**Motivation:** Enhancer-derived RNAs (eRNAs) and their fusion with protein-coding genes represent a crucial yet understudied layer of transcriptional regulation. eRNAs are typically expressed at low levels, which makes fusion events difficult to detect with conventional fusion detection tools. In addition, these tools are not designed to capture fusion transcripts arising from spatial proximity between distal regulatory elements and gene loci. Reads spanning such regions are also frequently filtered as mapping artifacts. As a result, computational approaches for systematically identifying spatially mediated enhancer–exon fusion transcripts remain lacking.

**Methods:** We developed ChiMER, a graph-based framework for detecting **ChiM**eric **E**nhancer **R**NAs from short-read RNA-seq data. ChiMER constructs splice graphs with chromatin contact information to introduce enhancer–exon edges and uses graph alignment to search for potential transcriptional paths. A ranking-based scoring module then prioritizes high-confidence events. Evaluations on simulated and real RNA-seq datasets show that ChiMER achieves higher sensitivity than conventional linear fusion detection methods while maintaining low false-positive rates.

**Results:** Applied to cancer cell line RNA-seq datasets, ChiMER identified multiple enhancer–exon chimeric transcripts, several associated with super-enhancer regions. Multi-omics analysis further shows that fusion transcripts occur in transcriptionally active regulatory environments and frequently coincide with strong R-loop signals, suggesting a potential role of RNA–DNA hybrid structures in facilitating long-range transcriptional joining events.

**Availability:** https://github.com/Candlelight-XYJ/ChiMER

## 1 Introduction

Enhancer RNAs (eRNAs) function as pivotal regulators of gene expression and exhibit divergent functional roles determined by their genomic origins. While intragenic eRNAs frequently coordinate with host-gene splicing machinery, intergenic eRNAs primarily serve as structural scaffolds that stabilize long-range chromatin loops (Li, et al., 2016; Sartorelli and Lauberth, 2020).

Fusion transcripts, defined as transcripts resulting from the fusion of different parent genes, are essential to biological processes (Barresi, et al., 2019; Dorney, et al., 2023). The formation of fusion transcripts generally occurs through two primary mechanisms. The first involves DNA recombination that generates fusion genes, which are subsequently transcribed into chimeric RNAs. This pathway is extensively documented for its role in the development of various cancers. The second mechanism involves RNA-level fusion, which includes transcriptional read-through and transsplicing. Due to the inherent complexity of these processes, significant gaps remain in the current understanding of RNA-level fusions. Building upon these classical mechanisms, researchers proposed the RNA-poise hypothesis in 2019 to describe the necessary conditions for the production of chimeric transcripts (Yan, et al., 2019). This hypothesis posits that the two participating RNA molecules must maintain spatial proximity. In 2021, researchers demonstrated that bidirectional transcription products from double-stranded DNA can form chimeric transcripts, a finding that reinforces the principle of spatial proximity suggested by the RNA-poise model (Wang, et al., 2021).

Previous research indicates that eRNAs are also capable of forming chimeric transcripts (Kowalczyk, et al., 2012). Specifically, eRNAs transcribed from intragenic enhancers can join with the exons of their host genes to form a novel class of chimeric RNAs known as meRNAs. These meRNAs possess polyA tails, which contribute to their stability and facilitate their significant regulatory functions within the cell.

Despite these findings, research on eRNA-derived chimeric RNAs remains scarce due to substantial computational and biological hurdles. First, systematic identification is hindered by current bioinformatics pipelines that automatically filter out enhancer annotations overlapping with known genes. This standard practice causes chimeric RNAs formed between eRNAs and other molecules to be discarded as computational noise (Haas, et al., 2019). Second, the low abundance of eRNAs relative to protein-coding transcripts makes it difficult to distinguish genuine fusion events from random transcriptional read-throughs in short-read sequencing data. Furthermore, conventional views suggest that intergenic enhancers serve exclusively as regulatory elements (Panigrahi and O’Malley, 2021). Consequently, whether eRNAs from intergenic enhancers can span chromatin loops to form fusion transcripts with their target mRNAs remains a critical yet unexplored question.

To address these limitations, we introduce ChiMER for the identification of chimeric enhancer RNAs from short-read RNA-seq data. ChiMER utilizes a specialized splice graph that incorporates exon-enhancer and exon-exon junctions as prior knowledge to constrain the search space. This approach enables the multi-step computational pipeline to identify fusion transcripts formed by enhancer RNAs and protein-coding genes with high precision. Multi-omics validation demonstrates the reliability of the formation of fusion transcripts by eRNAs across spatial distances, while benchmarking results indicate that ChiMER excels at recovering junction fusion transcripts that are typically missed by general-purpose fusion detection software. Ultimately, ChiMER provides a tool to explore the role of eRNAs in spatial gene regulation and offers a new perspective on the complexity of the human transcriptome.

## 2 Methods

### 2.1 Problem formulation

We formulate enhancer–exon fusion detection as a graph-constrained alignment inference problem. Let *G* = (*V, E*) denote a directed splice graph, where vertices *v* ∈ *V* represent genomic elements, including annotated exons and candidate enhancer regions, and edges *e* ∈ *E* represent transcriptionally permissible transitions. The edge set *E* encompasses canonical exon–exon junctions derived from gene annotations and novel enhancer–exon links inferred from regulatory priors.

Given a set of RNA sequencing reads *R*, the objective is to identify read-supported paths in *G* that traverse at least one enhancer–exon edge. Each read *r* ∈ *R* is aligned to the graph to yield an alignment path *Pr* = (*v*_1_, *v*_2_, …, *v*_*k*_), where consecutive vertices are connected by edges in *E*. A candidate enhancer–exon fusion event is identified if a path satisfies the following criteria: (i) the path contains at least one enhancer vertex and one exon vertex; (ii) the path includes at least one enhancer–exon edge; (iii) the alignment score exceeds a predefined threshold **τ**. The detection task thus reduces to identifying and aggregating all graph-supported alignment paths that satisfy these structural constraints.

### 2.2 ChiMER computational framework

#### 2.2.1 Construction of an enhancer-aware splice graph

ChiMER constructs a directed splice graph that integrates canonical transcript structures with regulatory spatial priors. Vertices in the graph represent two classes of genomic elements: (i) annotated protein-coding exons derived from reference gene annotations and (ii) candidate enhancer regions curated from public enhancer catalogs (Dyer, et al., 2025; Kawaji, et al., 2017). Each vertex is uniquely indexed and retains genomic coordinates along with strand information where available.

Canonical exon-exon edges are derived from transcript models by connecting consecutive exons within the same transcript, thereby preserving the annotated splicing structure. These edges define the baseline transcriptional backbone of the graph.

To incorporate regulatory information, enhancer-exon edges are introduced through two complementary mechanisms. First, the intragenic spatial edges link enhancers to the exons of the same gene when they locate on the same chromosome and within a predefined genomic distance window. Second, intergenic or distal edges are added based on enhancer-promoter interaction (EPI) information obtained from eRNAbase, which integrates evidence from both experimental assays and computational predictions (Song, et al., 2024). These edges reflect three-dimensional genomic proximity and are established if either locus overlaps with interaction anchors.

#### 2.2.2 Graph-constrained alignment and fusion inference

ChiMER employs a two-stage alignment strategy to identify potential enhancer-exon fusion events. In the first stage, short reads from RNA-seq data are aligned to the linear reference genome to extract candidate reads that generate split alignments. This step facilitates the localization of reads spanning two discontinuous genomic positions, thereby providing initial evidence for chimeric transcription. To minimize alignment ambiguity, only uniquely mapped reads are retained, and their breakpoint information is recorded for downstream processing.

Subsequently, these candidate split reads are aligned against a pre-constructed enhancer-aware splice graph (Rautiainen and Marschall, 2020). Unlike conventional linear genome alignment, graph-based alignment allows reads to traverse multiple edge types within the graph, including canonical exon-exon junctions and enhancer-exon links. This approach enables the evaluation of whether a read spans regulatory edges within a unified computational framework.

For each read, the graph alignment process generates a corresponding path representation. A read is identified as supporting a candidate fusion if its path includes at least one enhancer vertex and one exon vertex, traverses an enhancer-exon edge, and yields an alignment score exceeding a predefined threshold. By performing a constrained search within the graph space, ChiMER reformulates fusion detection as a path inference problem under structural priors, which effectively reduces false positive signals arising from repetitive sequences in linear alignments.

#### 2.2.3 Consensus sequence reconstruction

To enhance the reliability of fusion events, ChiMER introduces a read clustering and consensus sequence reconstruction stage following the graph-constrained alignment. Let a candidate enhancer-exon path *e* be supported by a set of reads *R*_*e*_ = {*r*_1_, *r*_2_, …, *r*_*n*_}. Initially, *R*_*e*_ is partitioned based on breakpoint positions and path structures within the graph space to ensure that reads within the same cluster correspond to an identical enhancer-exon junction pattern.

Within each read cluster, sequence fragments spanning the breakpoint region undergo multiple sequence integration to construct a local consensus sequence *C*_*e*_. This process aims to eliminate sequencing errors or local mismatches in individual reads, thereby yielding a more stable breakpoint representation. The resulting consensus sequence is re-evaluated for consistency with the graph path, and a consistency support ratio is calculated. To further minimize false positives, ChiMER imposes multiple filtering criteria on candidate events, including:(i) a minimum threshold for the number of supporting reads; (ii) consistency requirements between the consensus sequence and the graph path; (iii) the removal of alignments potentially originating from repetitive regions.

A candidate event is retained as a final fusion result only when it satisfies preset conditions for both read count and sequence consistency. By incorporating consensus reconstruction and structural consistency filtering, ChiMER elevates fusion identification from single-read evidence to cluster-level structural support, thereby enhancing the robustness of the results.

#### 2.2.4 Prioritization of candidate junctions

To assess the reliability of candidate enhancer-exon junctions, we first extract functional genomics features from their proximal regions, including H3K27ac, ATAC-seq, and CAGE-seq signals. Candidate junctions identified by ChiMER serve as positive samples, whereas non-promoter-enhancer interacting regions randomly sampled from the genome represent the negative samples. We define a linear scoring function to calculate a composite score for each sample:

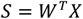

where *X* is the feature vector and *W* represents the learnable feature weights. The weights are optimized by minimizing a ranking loss function to ensure that scores for positive samples are consistently higher than those for negative samples. Specifically, we employ a ranking loss in the form of a hinge loss:

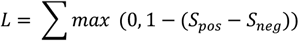

where *S*_*pos*_ and *S*_*neg*_ represent the scores of positive and negative samples, respectively. After determining the optimal weights, we compute scores for all candidate junctions. To evaluate statistical significance, we treat the distribution of negative sample scores as the null distribution and calculate an empirical p-value for each candidate:

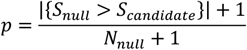

The candidate enhancer-exon junctions are ultimately ranked and filtered based on their scores and the associated significance levels.

### 2.3 Cross-omics data validation

To systematically evaluate the biological plausibility of the enhancer-exon fusion events identified by ChiMER, we establish a multi-omics validation framework. The validation datasets derived from the A549 and K562 cell lines, include ATAC-seq, CAGE-seq, H3K27ac ChIP-seq, HiC, and long-read sequencing data.

#### Validation of regulatory activity evidence

We first verify the regulatory activity at the enhancer. We perform an overlap analysis between the genomic positions of enhancers and genes within the predicted chimeric eRNAs and the peaks derived from ATAC-seq and H3K27ac ChIP-seq. A chimeric eRNA is considered to have chromatin-level support only if it significantly overlaps with open chromatin regions. Concurrently, CAGE-seq peaks are utilized to verify the presence of transcription initiation signals within the enhancer regions to assess their transcriptional activity.

#### Spatial interaction support

To verify the spatial proximity between the enhancer and the gene containing the target exon, we intersect the predicted enhancer-gene pairs with statistically significant Hi-C interaction intervals. If a significant chromatin interaction signal exists between the enhancer and the corresponding gene body, the fusion event is deemed to have structural support at the three-dimensional genomic level.

#### Structural validation via long-read sequencing

To further confirm the authenticity of the fusion structures, we extract split reads from long-read sequencing data that span the predicted breakpoint regions. Comparative alignment analysis is conducted to confirm whether the long reads contiguously cover both the enhancer and exon regions. The presence of long-read support directly bridging the two regions defines the event as passing structural-level validation.

#### Evidence integration strategy

For each candidate fusion event, we quantify the number of supporting multi-omics evidence types, including chromatin activity, spatial interaction, and long-read structural support. Events are stratified based on the count of these supporting types to evaluate the biological consistency of the ChiMER predictions.

### 2.4 Benchmarking settings

To systematically evaluate the detection performance of ChiMER, simulated datasets are generated using the polyester R package (Frazee, et al., 2015). The simulation design includes both negative and positive control groups. Negative samples consist exclusively of randomly selected expression signals from normal transcripts to assess false positive rates. Positive samples incorporate artificially constructed enhancer-exon fusion transcripts integrated into a background of normal transcript expression. Fusion sites are derived from enhancer-gene interaction pairs provided by the eRNAbase. These fusion transcripts are generated by concatenating 300 bp sequences from the respective upstream and downstream nodes, with the actual junctions recorded as the ground truth.

The simulation utilizes a multi-dimensional parameter matrix, including three random seeds, varying numbers of fusion events (10, 20, and 50), different fusion expression levels (10, 20, and 30), and multiple background expression levels (10, 30, and 60). The sequencing parameters are set to 150 bp paired-end reads. By systematically traversing this parameter matrix, scenarios with diverse signal-to-noise ratios are constructed.

Arriba and STAR-Fusion are selected as the baseline method for comparison and are executed under identical reference genome and annotation conditions (Haas, et al., 2017; Uhrig, et al., 2021). Recall, Precision, and F1-score are calculated based on the consistency between predicted and ground-truth breakpoints. Furthermore, the number of false positives is quantified in negative samples to evaluate the performance of different methods across various levels of complexity.

## 3 Results

### 3.1 Cross-omics validation of chimeric enhancer RNAs

The application of ChiMER to polyA RNA-seq data from the A549 and K562 cell lines identified 24 and 37 candidate enhancer-exon fusion transcripts, respectively (Supplementary Table 3 and Table 4). To verify the regulatory activity of these candidates, we integrate ATAC-seq, H3K27ac ChIP-seq, and CAGE-seq data. As illustrated in Figure 2, significant ATAC-seq and H3K27ac peaks are present at both the enhancer and the corresponding FYB1 gene body regions, indicating an open and active chromatin state. The enhancer region also exhibits distinct CAGE-seq signals, which support its transcriptional activity.

**Figure 1.**
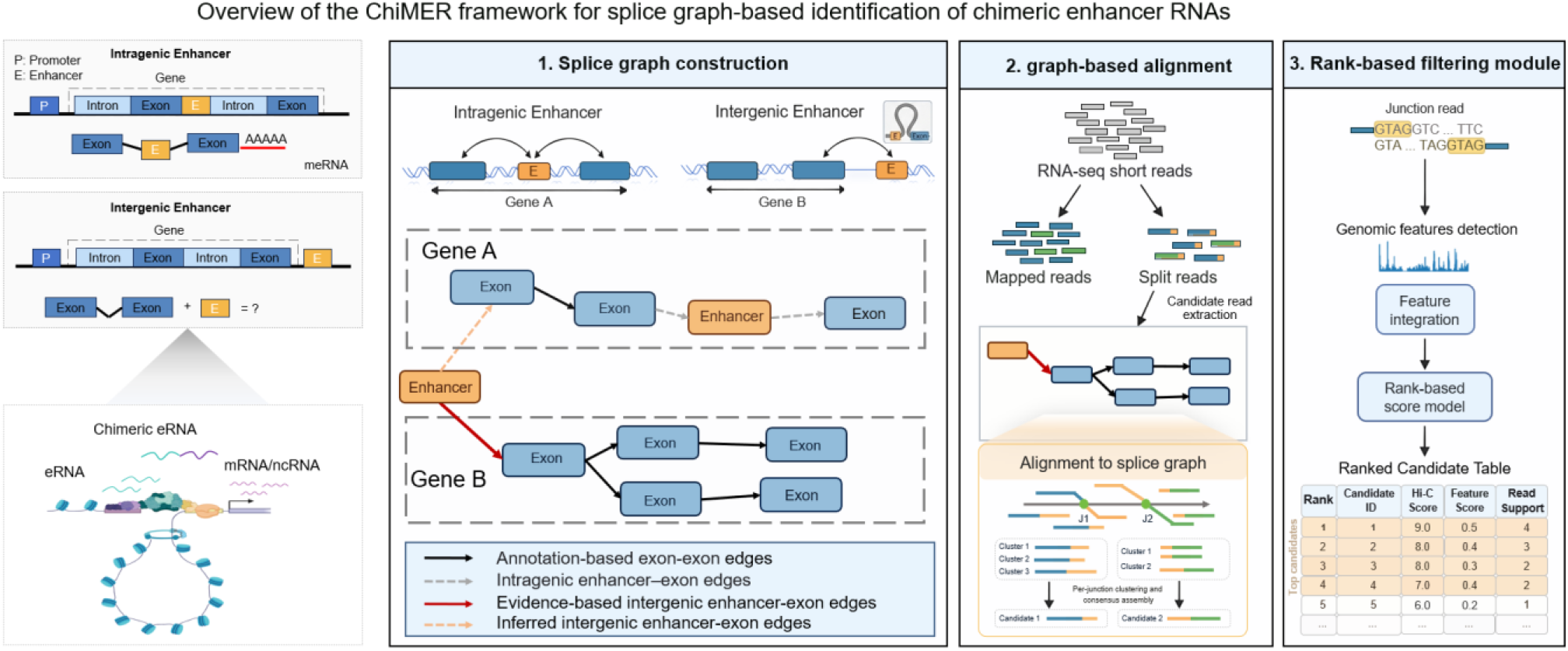
Overview of the ChiMER framework. Schematic overview of the ChiMER framework for detecting enhancer-exon chimeric transcripts. The pipeline consists of three primary modules: Step 1: Splice graph construction, where a comprehensive genomic graph is built by integrating canonical exon-exon junctions with intergenic or intragenic enhancer-exon prior edges. Step 2: Graph-based alignment, which involves mapping RNA-seq split reads onto the constructed graph to identify potential chimeric junctions. Step 3: Scoring module, where candidate transcripts are prioritized and filtered based on evidence strength and statistical significance to ensure high-precision fusion detection.

**Figure 2.**
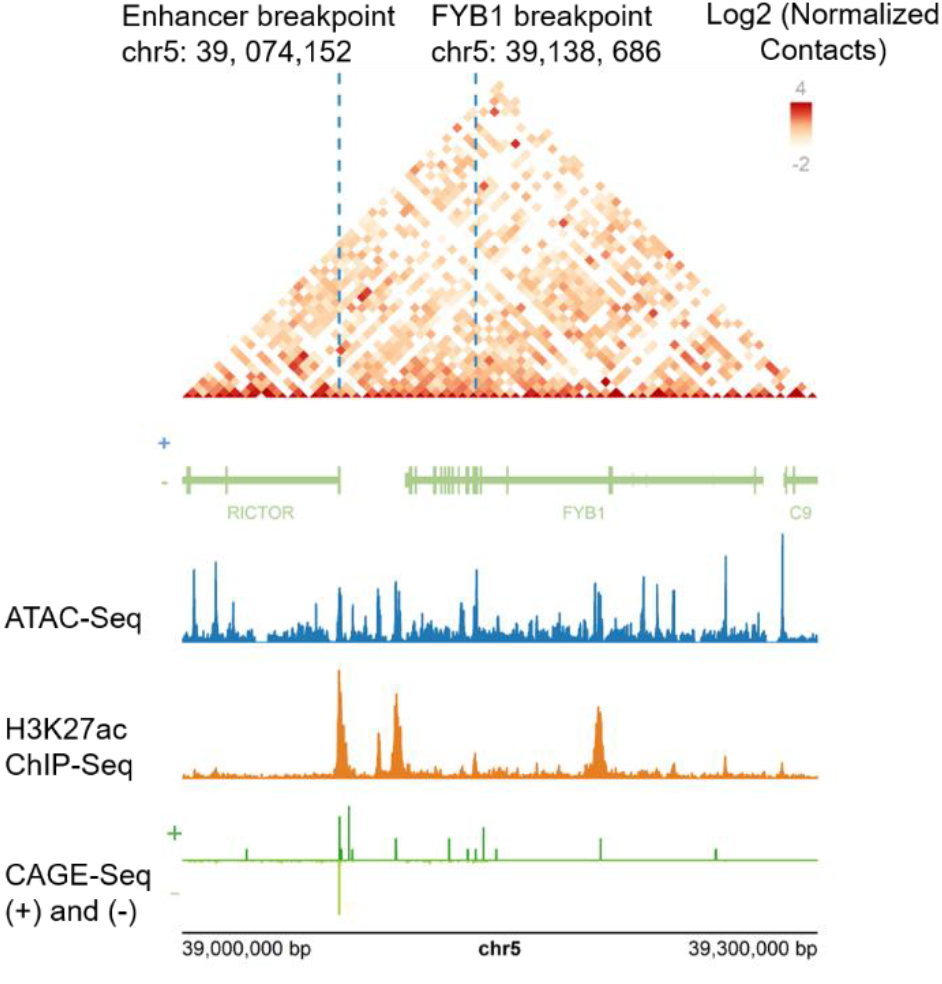
Spatial and transcriptional validation of chimeric eRNA fusion events in A549 cell line. Genome browser view showing ATAC-seq, H3K27ac ChIP-seq, and CAGE-seq signals at the predicted enhancer and exon loci. Enrichment of chromatin accessibility and active histone marks supports regulatory activity at both regions. Hi-C contact map illustrating spatial proximity between the enhancer and the target gene locus. The interaction occurs within the same TAD, indicating three-dimensional structural support. Genome version: GRCh38.

Further analysis of the spatial structural relationships using Hi-C data, as shown in Figure 2, reveals that the enhancer and the target gene are located within the same topologically associating domain (TAD) and display significant contact signals, suggesting spatial proximity within the three-dimensional genome. Similar multi-omics integration and spatial proximity were also observed for candidates in K562 cells, such as the SPRED2 (Supplementary Figure 1).

Long-read sequencing results demonstrate the presence of contiguous reads spanning the predicted junctions (Figure 3). The breakpoint positions in the long reads are consistent with the short-read data.

**Figure 3.**
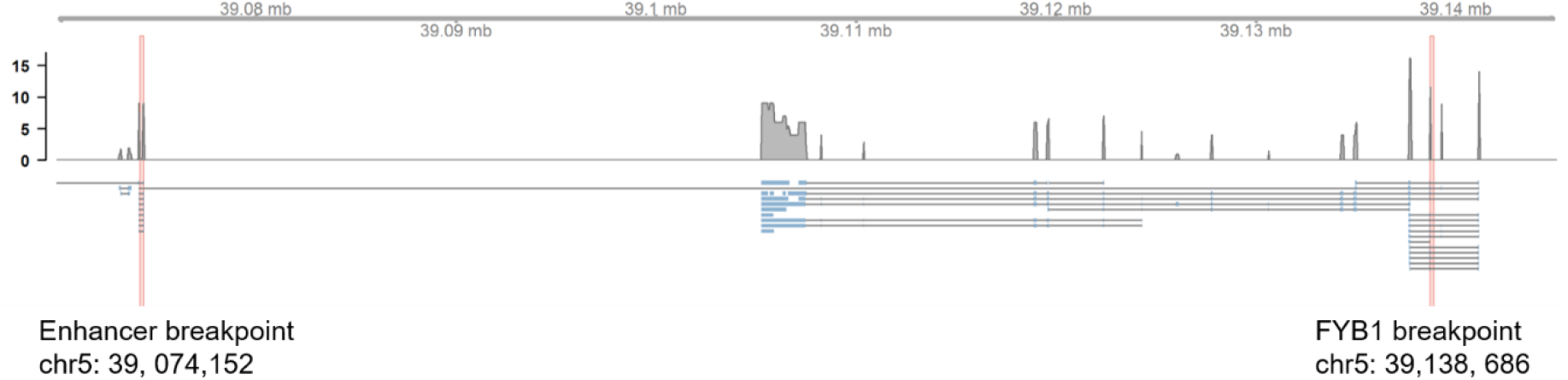
Long-read sequencing validation of the enhancer (chr5: 39, 074, 152) and FYB1 fusion event. Genomic visualization of Nanopore long-reads on chr5. The coverage track (top) and split-read alignment track (bottom) demonstrate the physical continuity of the fusion transcript. Red dashed lines mark the genomic breakpoints. Each blue-segmented line represents a single chimeric read bridging the gap between the enhancer breakpoint and FYB1 breakpoint.

FYB1 has previously been reported to be regulated by a super-enhancer in T-ALL cell lines, indicating its susceptibility to enhancer-driven transcriptional control (Zhang, et al., 2023). These collective results indicate that the enhancer-exon fusion events identified by ChiMER are biologically plausible in terms of both transcriptional activity and spatial organization.

### 3.2 Performance evaluation on simulated datasets

To systematically evaluate the performance of ChiMER, we design two sets of simulation experiments consisting of positive and negative controls.

The positive control is primarily used to evaluate the ability of the tools to accurately identify enhancer-exon fusion events from complex background transcripts under various signal-to-noise ratios. This experiment constructs multiple complexity scenarios by controlling the number of fusion events, fusion expression levels, and background expression intensity. As illustrated in Table 1, despite low fusion expression and high background noise, the F1-score of ChiMER is higher than Arriba and STAR-Fusion. ChiMER also exhibits a more stable performance trend across different parameter combinations (Supplementary Figure 2).

**Table 1.** F1-score of ChiMER and Arriba in detecting enhancer-exon fusions under simulated conditions.

| Dataset | Background transcript abundance | Fusion transcript abundance | ChiMER | Arriba | STAR-Fusion |
| --- | --- | --- | --- | --- | --- |
| Positive | 10 | 10 | $0.628 \pm 0.101$ | 0.00 | 0.00 |
| Positive | 30 | 10 | $0.628 \pm 0.055$ | 0.00 | 0.00 |
| Positive | 60 | 10 | $0.593 \pm 0.058$ | 0.00 | 0.00 |
| Negative | 10 | 0 | 1.00 | 1.00 | 1.00 |
| Negative | 30 | 0 | 1.00 | 1.00 | 1.00 |
| Negative | 60 | 0 | 1.00 | 1.00 | 1.00 |

Simulations were performed using Polyester (n=3 random seeds). Fusion and background transcript abundances were controlled by the reads\_per\_transcript parameter. The number of ground-truth fusion events was fixed at 20.

The negative control is primarily used to assess the capacity for false positive control. Negative samples consist exclusively of expression signals from normal transcripts and do not contain any artificially constructed fusion events. As shown in Table 1, across all background expression levels, the number of fusions predicted by ChiMER is 0, which is equal to Arriba and STAR-Fusion. This indicates that the graph-structure constraint strategy facilitates the reduction of false detections caused by random splicing.

Overall, the simulation data indicate that ChiMER enhances the detection capability for low-expression fusion events while maintaining high precision and effectively controlling false positive levels.

### 3.3 Effects of graph prior on candidate detection

We evaluated the influence of graph priors on candidate detection using the A549 RNA-seq dataset by comparing two configurations: a linear splice graph and a full spatial splice graph. The linear graph contains only canonical exon–exon edges, while the full spatial graph incorporates all enhancer–exon interactions derived from spatial data.

As shown in Table 2, the number of edges increases from 408285 (linear) to 18846931 (full graph), accompanied by an increase in split-read supported junctions from 0 and 2932. Consequently, the full graph identifies 24 candidate transcripts, compared with 0 in the linear graph, indicating that spatially informed priors improve the detection of enhancer–exon transcripts beyond canonical splicing.

**Table 2.** Effect of graph priors on candidate detection in the A549 dataset.

| Graph prior | No. Edges | No. Split reads after graph alignment | No. Candidates |
| --- | --- | --- | --- |
| Full graph | 18846931 | 2932 | 24 |
| Linear graph | 408285 | 0 | 0 |

### 3.4 Pattern analysis of intragenic and intergenic chimeric enhancer RNAs

As illustrated in Figure 4A, the genomic distances for intragenic and intergenic enhancer-exon fusions exhibit distinct distribution characteristics. The distances for intragenic events are primarily concentrated within the range of 1 kb to 10 kb. In contrast, the distribution of intergenic events is more dispersed, with a median distance of 100 kb and a maximum extension of up to 30 Mb. These findings suggest that while most fusion events occur within a relatively localized regulatory scope, long-range connectivity also exists.

**Figure 4.**
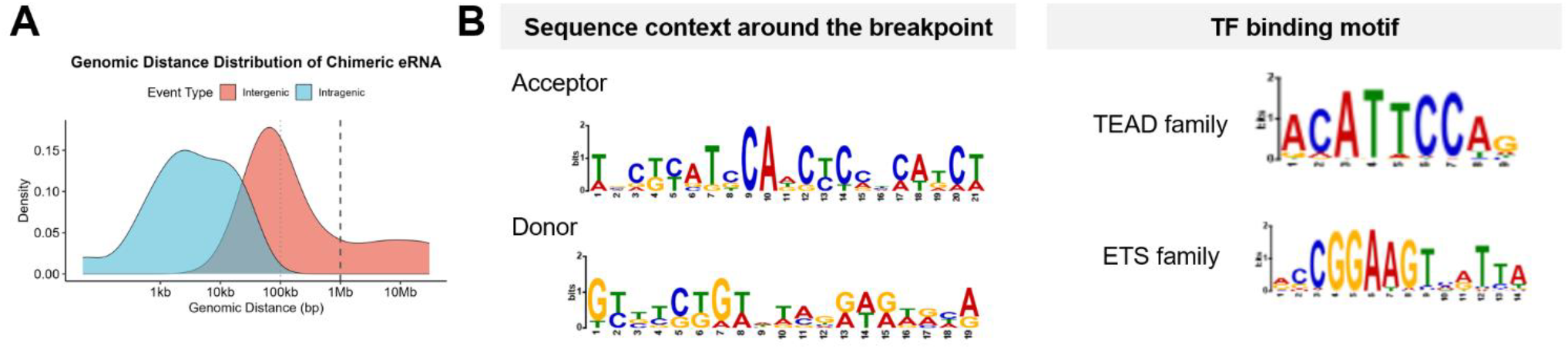
Genomic distance and motif analysis of chimeric eRNAs.

In terms of sequence characteristics at the breakpoints, we perform a motif analysis of the breakpoint site regions. As shown in Figure 4B, these intergenic chimeric eRNA breakpoints lack the characteristic GT/AG splice consensus sequences, suggesting that the observed junctions are unlikely to result from conventional splicing. To examine the surrounding sequence context, we expanded the genomic windows for motif enrichment analysis. Sequences spanning 100 bp upstream to 3 bp downstream of putative acceptor sites and 3 bp upstream to 100 bp downstream of donor sites were extracted and analyzed using the AME tool (McLeay and Bailey, 2010). The results show significant enrichment of several transcription factor binding motifs, particularly those from the TEAD and ETS families. These transcription factors are commonly associated with active regulatory elements and enhancer-related transcription, suggesting that the breakpoints occur in transcriptionally active regulatory regions rather than canonical splice junctions.

### 3.5 R-loop signals at breakpoint regions of enhancer–exon fusion transcripts

In the A549 dataset, ChiMER identifies a fusion transcript formed by a super-enhancer and its target gene FYB1. Similarly, the fusion transcript detected in the K562 cell line involves SPRED2 and its proximal super-enhancer. Super-enhancers typically exhibit a highly active transcriptional state and generate substantial amounts of enhancer RNA. Previous studies establish that active transcription and eRNA production within enhancer regions are closely associated with the formation of R-loops, which are DNA:RNA hybrid structures (Jia, et al., 2023; Watts, et al., 2022). Based on DRIP-seq data from the K562 cell line (Figure 5), distinct R-loop signal peaks are observed near the two breakpoints of SPRED2 and the associated super-enhancer. These results indicate that the fusion transcript breakpoints are likely situated within regulatory regions characterized by high transcriptional activity and concomitant R-loop formation. During transcription, R-loops can alter local DNA architecture and influence RNA polymerase initiation, elongation, and termination processes. Such structures may create favorable conditions for non-canonical RNA ligation events, thereby facilitating the production of enhancer RNA-exon chimeric transcripts.

**Figure 5.**
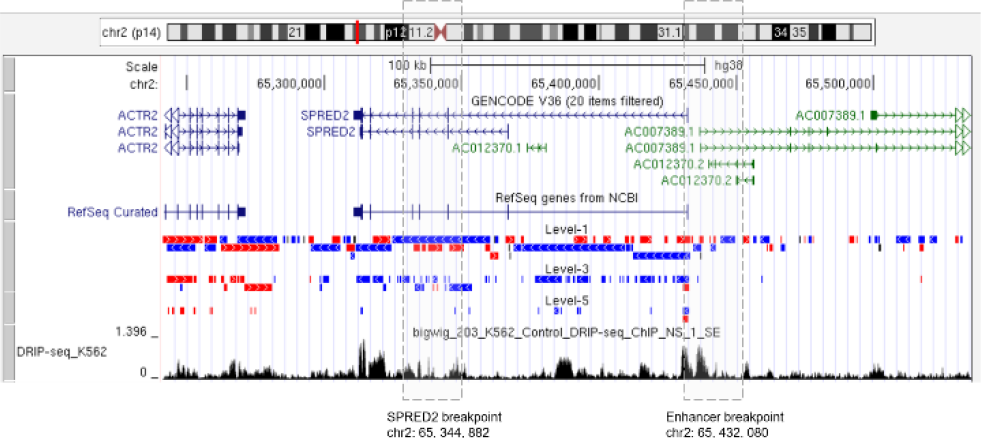
R-loop signals at breakpoint regions in K562 cells. Both breakpoint loci overlap with peaks of R-loop signal, suggesting that these regions reside within transcriptionally active loci where RNA– DNA hybrid structures are present.

## 4 Discussion

Current fusion transcript detection tools are primarily designed for canonical transcripts. Their filtering strategies often assume that non-canonical breakpoint signals predominantly originate from sequencing noise or alignment errors. Consequently, fusion signals located in enhancer regions are frequently discarded as low-confidence events. However, several studies demonstrate that enhancers themselves possess transcriptional activity and generate enhancer RNAs (eRNAs). Whether eRNA-involved fusions represent a genuine regulatory layer remains unexplored. The ChiMER framework proposed in this study attempts to systematically mine these potential transcriptional dark matter signals beyond conventional noise filtering.

Methodologically, ChiMER employs graph-structure constraints to restrict candidate breakpoints to biologically interpretable splicing paths. In simulations, this targeted filtering effectively suppresses background noise and false positives, thereby enhancing the detection of low-expression fusion events. Multi-omics support in real datasets further suggests that these enhancer-exon fusions are not random ligations but are likely closely associated with enhancer activity states and genomic architecture.

From a mechanistic perspective, the formation of chimeric eRNAs may involve several pathways. First, enhancer regions themselves exhibit bidirectional transcriptional activity. When spatially proximal to neighboring genes, continuous transcripts may be generated through transcriptional extension or read-through transcription. Second, spatial contacts between enhancers and genes may create conditions that favor non-canonical splicing. In particular, enhancer transcription can promote the formation of RNA– DNA hybrid structures, which may facilitate splice events between genomic regions that are spatially proximal but linearly distant. Chromatin remodeling and regulatory imbalance in cancer cells may further increase the likelihood of such events. Although this study cannot directly prove specific molecular mechanisms, consistent multi-omics evidence supports their potential biological foundation.

It should be noted that the construction of the ChiMER splice graph relies on existing annotations and spatial interaction data, which may not cover all potential connection forms. While using fusion events derived from graph-structure edge sets in simulations helps evaluate performance in targeted scenarios, future studies are required to assess the generalization capability of the method using independently constructed random connection models or additional experimental validation data. Moreover, this study is primarily based on the analysis of A549 and K562 cell lines; the prevalence and functional significance of enhancer-exon fusions across different tissue types and disease contexts remain to be investigated in depth.

Overall, ChiMER provides a new strategy for the systematic exploration of enhancer-related fusion transcripts. By mining inter-regional transcriptional signals neglected by traditional analytical pipelines, this study offers a new research direction for understanding the potential links between enhancer transcriptional activity and gene expression regulation.

## Acknowledgements

None.

