## Supplementary Information for "ChiMER: Integrating chromatin architecture into splicing graphs for chimeric enhancer RNAs detection"

**Implementation details and data preprocessing**

The human GRCh38 (hg38) genome sequence and corresponding gene annotation files were downloaded from the Ensembl genome browser (Ensembl release 110). Enhancer coordinates, eRNA annotations, and enhancer-promoter interactions specific to the A549 cell line are obtained from the eRNAbase database (http://www.licpathway.net/eRNAbase/index.php). Multi-omics validation datasets, comprising paired-end RNA-Seq, ATAC-Seq, H3K27ac ChIP-Seq, CAGE-Seq, and Hi-C data, are retrieved from the ENCODE Project portal (details are listed in Supplementary Table 1 and Table 2). Raw A549 Nanopore long-read RNA sequencing data for molecular structure validation are sourced from the NCBI SRA database. Simulation datasets for performance evaluation are generated via the Polyester tool, which simulates various expression levels and signal-to-noise conditions through the adjustment of the reads_per_transcript parameter. Quality control and adapter removal for all raw sequencing data are conducted using fastp. Algorithm performance is evaluated through a multilayered validation system. Benchmarking of fusion detection performance on simulated data is conducted using Arriba (v2.5.1) as a reference tool. Motif enrichment analysis is performed on high-confidence eRNA-exon fusion breakpoints using the MEME Suite.

The ChiMER algorithm employs a hierarchical alignment strategy to improve the accuracy of chimeric transcript identification. Initial linear genomic alignment is performed using STAR, where only uniquely mapped results are retained via the --outFilterMultimapNmax 1 parameter and chimeric alignment mode is enabled with a minimum segment length of 15 bp through --chimSegmentMin 15 to capture potential chimeric breakpoints. Chimeric junction information is output separately, while partially unmapped reads are preserved for further analysis. To resolve complex chimeric structures that are difficult to precisely characterize through linear alignment, ChiMER utilizes GraphAligner to map reads against a pre-constructed A549-specific splicing graph in GFA format. The graph alignment process sets the minimum seed length to 20 bp, the alignment score threshold to 0.5, and the bandwidth to 500, while allowing multimapping with a score fraction of 0.9 to balance sensitivity and specificity.

**Supplementary Table 1. Datasets used in this study for chimeric eRNAs identification**

| # GEO Accession | Assay Type | Sample | Project | Organism | SRR ID |
| --- | --- | --- | --- | --- | --- |
| GSM5379323 | polyA plus RNA-seq | A549 | ENCODE | Homo sapiens | SRR14843205 |
| GSM8672557 | polyA plus RNA-seq | K562 | ENCODE | Homo sapiens | SRR31643422 |

**Supplementary Table 2. Multi-omics data for validation**

| Omics Data | Sample | Project | Organism | Links/ID |
| --- | --- | --- | --- | --- |
| ATAC-Seq | A549 | ENCODE | Homo sapiens | <https://www.encodeproject.org/files/ENCFF795ASF/> |
| H3K27ac Chip-Seq | A549 | ENCODE | Homo sapiens | <https://www.encodeproject.org/files/ENCFF654HMM/> |
| CAGE-Seq | A549 | ENCODE | Homo sapiens | <https://www.encodeproject.org/files/ENCFF368QQU/> |
| Hi-C | A549 | ENCODE | Homo sapiens | https://www.encodeproject.org/experiments/ENCSR444WCZ/ |
| Long Read RNA-Seq | A549 | SRA | Homo sapiens | https://trace.ncbi.nlm.nih.gov/Traces/?run=SRR15045006 |
| ATAC-Seq | K562 | GEO | Homo sapiens | GSM6637785 |
| H3K27ac Chip-Seq | K562 | ENCODE | Homo sapiens | https://www.encodeproject.org/files/ENCFF532MMV/ |
| CAGE-Seq | K562 | ENCODE | Homo sapiens | https://www.encodeproject.org/files/ENCFF156SGB/ |
| Hi-C | K562 | 3D Genome | Homo sapiens | https://3dgenome.fsm.northwestern.edu/datasets?id=128 |
| Long Read RNA-Seq | K562 | GEO | Homo sapiens | GSM5331401 |
| DRIP-Seq | K562 | R-loopBase | Homo sapiens | <https://rloopbase.nju.edu.cn/> |

**Supplementary Table 3. Candidate chimeric eRNAs identified by ChiMER in A549 cell line**

| **chrA** | **breakpoint_posA** | **strandA** | **chrB** | **breakpoint_posB** | **strandB** | **enhancer_id** | **gene_id** | **hic_support** | **atac_overlap** | **chip_overlap** | **cage_overlap** | **support_reads** | **nano_support_count** | **in_same_tad** | **distance** | **type** |
| --- | --- | --- | --- | --- | --- | --- | --- | --- | --- | --- | --- | --- | --- | --- | --- | --- |
| chrX | 1.54E+08 | + | chrX | 154368087 | + | Enh_chrX_154367038_154367838 | ENSG00000196924 | TRUE | TRUE | TRUE | TRUE | 1 | 277 | TRUE | 2828 | Intragenic |
| chr8 | 61684183 | - | chr8 | 61714307 | - | Enh_chr8_61722287_61722887 | ENSG00000198363 | TRUE | TRUE | TRUE | TRUE | 1 | 1657 | TRUE | 30124 | Intergenic |
| chr5 | 39138686 | + | chr5 | 39074152 | + | Enh_chr5_39178141_39178941 | ENSG00000082074 | TRUE | TRUE | TRUE | TRUE | 8 | 9 | TRUE | 64534 | Intergenic |
| chr5 | 39141137 | + | chr5 | 39074154 | + | Enh_chr5_39021141_39021341 | ENSG00000164327 | TRUE | TRUE | TRUE | TRUE | 6 | 9 | TRUE | 66983 | Intergenic |
| chr4 | 57109902 | + | chr4 | 57040912 | + | Enh_chr4_57049277_57049877 | ENSG00000163453 | TRUE | TRUE | TRUE | TRUE | 2 | 402 | TRUE | 68990 | Intergenic |
| chr3 | 69080138 | + | chr3 | 69077901 | + | Enh_chr3_69200759_69201159 | ENSG00000144744 | TRUE | TRUE | TRUE | TRUE | 1 | 51 | TRUE | 2237 | Intragenic |
| chr2 | 1.06E+08 | + | chr2 | 106164767 | + | Enh_chr2_106147112_106147912 | ENSG00000115652 | TRUE | TRUE | TRUE | TRUE | 1 | 76 | TRUE | 29384 | Intragenic |
| chr16 | 57479540 | + | chr16 | 57475604 | + | Enh_chr16_57404987_57405787 | ENSG00000125170 | TRUE | TRUE | TRUE | TRUE | 1 | 66 | TRUE | 3936 | Intragenic |
| chr16 | 30096092 | - | chr16 | 30096142 | - | Enh_chr16_30093578_30093978 | ENSG00000090238 | TRUE | TRUE | TRUE | TRUE | 1 | 16 | TRUE | 50 | Intragenic |
| chr15 | 29800683 | - | chr15 | 29822022 | - | Enh_chr15_29800505_29800705 | ENSG00000104067 | TRUE | TRUE | TRUE | TRUE | 2 | 109 | TRUE | 21339 | Intragenic |
| chr14 | 74829329 | - | chr14 | 74712932 | + | Enh_chr14_74704944_74705544 | ENSG00000119682 | TRUE | TRUE | TRUE | TRUE | 1 | 2 | TRUE | 116397 | Intergenic |
| chr14 | 54441268 | - | chr14 | 54436429 | + | Enh_chr14_54436132_54436732 | ENSG00000100528 | TRUE | TRUE | TRUE | TRUE | 1 | 474 | TRUE | 4839 | Intragenic |
| chr14 | 23069606 | - | chr14 | 23071163 | - | Enh_chr14_23035551_23035751 | ENSG00000100813 | TRUE | TRUE | TRUE | TRUE | 1 | 66 | TRUE | 1557 | Intragenic |
| chr13 | 98519458 | - | chr13 | 98576780 | - | Enh_chr13_98480745_98482145 | ENSG00000102572 | TRUE | TRUE | TRUE | TRUE | 1 | 98 | TRUE | 57322 | Intergenic |
| chr13 | 24512733 | + | chr13 | 24503763 | + | Enh_chr13_24501862_24502262 | ENSG00000102699 | TRUE | TRUE | TRUE | TRUE | 1 | 38 | TRUE | 8970 | Intragenic |
| chr12 | 52898710 | + | chr12 | 52288006 | + | Enh_chr12_52903749_52903949 | ENSG00000170421 | TRUE | TRUE | TRUE | TRUE | 2 | 3 | TRUE | 610704 | Intergenic |
| chr12 | 52316879 | + | chr12 | 52288028 | + | Enh_chr12_52230549_52231949 | ENSG00000170523 | TRUE | TRUE | TRUE | TRUE | 1 | 209 | TRUE | 28851 | Intragenic |
| chr12 | 52233616 | - | chr12 | 52235199 | - | Enh_chr12_52211349_52211549 | ENSG00000135480 | TRUE | TRUE | TRUE | TRUE | 2 | 7010 | TRUE | 1583 | Intragenic |
| chr10 | 1.19E+08 | + | chr10 | 119177136 | + | Enh_chr10_119174898_119175098 | ENSG00000165672 | TRUE | TRUE | TRUE | TRUE | 2 | 1002 | TRUE | 1641 | Intragenic |
| chr1 | 15807604 | + | chr1 | 46303728 | - | Enh_chr1_46314941_46315541 | ENSG00000173660 | TRUE | TRUE | TRUE | TRUE | 31 | 1187 | TRUE | 30496124 | Intergenic |
| chr1 | 2.02E+08 | + | chr1 | 201496299 | + | Enh_chr1_201491249_201491649 | ENSG00000159176 | TRUE | TRUE | TRUE | TRUE | 3 | 460 | TRUE | 10808 | Intragenic |
| chr1 | 1.79E+08 | + | chr1 | 179131466 | + | Enh_chr1_179179242_179179842 | ENSG00000143322 | TRUE | TRUE | TRUE | TRUE | 1 | 13 | TRUE | 11502 | Intragenic |
| chr1 | 1.57E+08 | + | chr1 | 156569450 | + | Enh_chr1_156602184_156602584 | ENSG00000183856 | TRUE | TRUE | TRUE | TRUE | 1 | 14 | TRUE | 3071 | Intragenic |
| chr1 | 1.54E+08 | + | chr1 | 153545508 | + | Enh_chr1_153569700_153569900 | ENSG00000196154 | TRUE | TRUE | TRUE | TRUE | 6 | 8482 | TRUE | 278 | Intragenic |

**Supplementary Table 4. Candidate chimeric eRNAs identified by ChiMER in K562 cell line**

| **chrA** | **breakpoint_posA** | **strandA** | **chrB** | **breakpoint_posB** | **strandB** | **enhancer_id** | **gene_id** | **hic_support** | **atac_overlap** | **chip_overlap** | **cage_overlap** | **support_reads** | **nano_support_count** | **both_in_A** | **in_same_tad** | **dist_val** |
| --- | --- | --- | --- | --- | --- | --- | --- | --- | --- | --- | --- | --- | --- | --- | --- | --- |
| chr5 | 54310476 | + | chr5 | 54171844 | + | Enh_chr5_54292438_54330115 | ENSG00000185305 | TRUE | TRUE | TRUE | TRUE | 3 | 80 | TRUE | TRUE | 138632 |
| chr5 | 1.27E+08 | + | chr5 | 1.27E+08 | + | Enh_chr5_126970143_126983862 | ENSG00000173926 | TRUE | TRUE | TRUE | TRUE | 4 | 49 | TRUE | TRUE | 112216 |
| chr2 | 65432080 | + | chr2 | 65344882 | + | Enh_chr2_65421149_65438694 | ENSG00000198369 | TRUE | TRUE | TRUE | TRUE | 6 | 15 | TRUE | TRUE | 87198 |
| chr7 | 43729488 | + | chr7 | 43648605 | + | Enh_chr7_43726465_43779451 | ENSG00000106603 | TRUE | TRUE | TRUE | TRUE | 11 | 134 | TRUE | TRUE | 80883 |
| chr13 | 41060881 | + | chr13 | 40982261 | + | Enh_chr13_40974343_41020895 | ENSG00000120690 | TRUE | TRUE | TRUE | TRUE | 1 | 32 | TRUE | TRUE | 78620 |
| chr10 | 22003311 | + | chr10 | 21929090 | + | Enh_chr10_23037598_23068341 | ENSG00000136770 | TRUE | TRUE | TRUE | TRUE | 1 | 115 | TRUE | TRUE | 74221 |
| chr10 | 22003265 | + | chr10 | 21929056 | + | Enh_chr10_23037598_23068341 | ENSG00000136770 | TRUE | TRUE | TRUE | TRUE | 6 | 115 | TRUE | TRUE | 74209 |
| chr4 | 1.03E+08 | + | chr4 | 1.03E+08 | + | Enh_chr4_102824422_102829675 | ENSG00000109332 | TRUE | TRUE | TRUE | TRUE | 2 | 19 | TRUE | TRUE | 58969 |
| chr1 | 2.47E+08 | - | chr1 | 2.47E+08 | - | Enh_chr1_246568274_246588073 | ENSG00000162852 | TRUE | TRUE | TRUE | TRUE | 1 | 154 | TRUE | TRUE | 24900 |
| chr3 | 9397878 | - | chr3 | 9422775 | - | Enh_chr3_9395173_9403218 | ENSG00000168137 | TRUE | TRUE | TRUE | TRUE | 1 | 16 | TRUE | TRUE | 24897 |
| chr7 | 1.21E+08 | - | chr7 | 1.21E+08 | - | Enh_chr7_121134128_121169788 | ENSG00000106034 | TRUE | TRUE | TRUE | TRUE | 11 | 564 | TRUE | TRUE | 21164 |
| chr12 | 31729028 | + | chr12 | 31709345 | + | Enh_chr12_31714270_31751415 | ENSG00000151743 | TRUE | TRUE | TRUE | TRUE | 2 | 11 | TRUE | TRUE | 19683 |
| chr10 | 71851236 | + | chr10 | 71834447 | + | Enh_chr10_71836616_71889797 | ENSG00000197746 | TRUE | TRUE | TRUE | TRUE | 6 | 1186 | TRUE | TRUE | 16789 |
| chr2 | 38602444 | + | chr2 | 38585863 | + | Enh_chr2_38593663_38671085 | ENSG00000143889 | TRUE | TRUE | TRUE | TRUE | 1 | 338 | TRUE | TRUE | 16581 |
| chr7 | 32891588 | + | chr7 | 32879844 | + | Enh_chr7_32877781_32906998 | ENSG00000170852 | TRUE | TRUE | TRUE | TRUE | 1 | 183 | TRUE | TRUE | 11744 |
| chr12 | 54391298 | + | chr12 | 54384489 | + | Enh_chr12_54351248_54394316 | ENSG00000161642 | TRUE | TRUE | TRUE | TRUE | 6 | 1 | TRUE | TRUE | 6809 |
| chr1 | 1.57E+08 | - | chr1 | 1.57E+08 | - | Enh_chr1_156745907_156773793 | ENSG00000143321 | TRUE | TRUE | TRUE | TRUE | 122 | 1171 | TRUE | TRUE | 6019 |
| chr12 | 57078695 | + | chr12 | 57072839 | + | Enh_chr12_57069711_57103675 | ENSG00000166881 | TRUE | TRUE | TRUE | TRUE | 1 | 15 | TRUE | TRUE | 5856 |
| chr12 | 54280051 | + | chr12 | 54274318 | + | Enh_chr12_54277166_54308323 | ENSG00000094916 | TRUE | TRUE | TRUE | TRUE | 3 | 71 | TRUE | TRUE | 5733 |
| chr7 | 1.03E+08 | + | chr7 | 1.03E+08 | + | Enh_chr7_102970978_102993961 | ENSG00000161040 | TRUE | TRUE | TRUE | TRUE | 14 | 115 | TRUE | TRUE | 5175 |
| chr6 | 30684510 | + | chr6 | 30679342 | + | Enh_chr6_30679761_30688902 | ENSG00000146112 | TRUE | TRUE | TRUE | TRUE | 15 | 54 | TRUE | TRUE | 5168 |
| chr1 | 1.6E+08 | + | chr1 | 1.6E+08 | + | Enh_chr1_159910036_159926488 | ENSG00000158710 | TRUE | TRUE | TRUE | TRUE | 1 | 615 | TRUE | TRUE | 5045 |
| chr10 | 72216050 | - | chr10 | 72220176 | - | Enh_chr10_72290628_72328614 | ENSG00000166295 | TRUE | TRUE | TRUE | TRUE | 1 | 200 | TRUE | TRUE | 4126 |
| chr1 | 8874880 | - | chr1 | 8878673 | - | Enh_chr1_8870761_8881067 | ENSG00000074800 | TRUE | TRUE | TRUE | TRUE | 257 | 9209 | TRUE | TRUE | 3793 |
| chr5 | 1.39E+08 | + | chr5 | 1.39E+08 | + | Enh_chr5_139468338_139496335 | ENSG00000184584 | TRUE | TRUE | TRUE | TRUE | 1 | 81 | TRUE | TRUE | 3790 |
| chr1 | 8878640 | + | chr1 | 8874913 | + | Enh_chr1_8870761_8881067 | ENSG00000074800 | TRUE | TRUE | TRUE | TRUE | 19 | 9209 | TRUE | TRUE | 3727 |
| chr11 | 65900385 | + | chr11 | 65897001 | + | Enh_chr11_65889780_65912542 | ENSG00000175592 | TRUE | TRUE | TRUE | TRUE | 1 | 39 | TRUE | TRUE | 3384 |
| chr12 | 57111216 | + | chr12 | 57108287 | + | Enh_chr12_57069711_57103675 | ENSG00000166888 | TRUE | TRUE | TRUE | TRUE | 3 | 13 | TRUE | TRUE | 2929 |
| chr6 | 43629016 | + | chr6 | 43626333 | + | Enh_chr6_43608487_43640618 | ENSG00000172432 | TRUE | TRUE | TRUE | TRUE | 2 | 28 | TRUE | TRUE | 2683 |
| chr2 | 15561206 | + | chr2 | 15558579 | + | Enh_chr2_15553698_15563705 | ENSG00000151779 | TRUE | TRUE | TRUE | TRUE | 1 | 15 | TRUE | TRUE | 2627 |
| chr8 | 67061919 | + | chr8 | 67059413 | + | Enh_chr8_67054444_67066837 | ENSG00000121022 | TRUE | TRUE | TRUE | TRUE | 3 | 259 | TRUE | TRUE | 2506 |
| chr1 | 1.13E+08 | + | chr1 | 1.13E+08 | + | Enh_chr1_112697982_112707738 | ENSG00000007341 | TRUE | TRUE | TRUE | TRUE | 1 | 35 | TRUE | TRUE | 2389 |
| chr1 | 1.51E+08 | + | chr1 | 1.51E+08 | + | Enh_chr1_151051179_151065229 | ENSG00000197622 | TRUE | TRUE | TRUE | TRUE | 1 | 344 | TRUE | TRUE | 1627 |
| chr1 | 23800753 | + | chr1 | 23799365 | + | Enh_chr1_23776065_23795725 | ENSG00000117308 | TRUE | TRUE | TRUE | TRUE | 3 | 355 | TRUE | TRUE | 1388 |
| chr1 | 2.02E+08 | + | chr1 | 2.02E+08 | + | Enh_chr1_202157528_202167214 | ENSG00000143851 | TRUE | TRUE | TRUE | TRUE | 3 | 152 | TRUE | TRUE | 1231 |
| chr11 | 62671307 | + | chr11 | 62670130 | + | Enh_chr11_62663535_62674099 | ENSG00000162194 | TRUE | TRUE | TRUE | TRUE | 1 | 178 | TRUE | TRUE | 1177 |
| chr11 | 1.19E+08 | - | chr11 | 1.19E+08 | - | Enh_chr11_119083976_119092619 | ENSG00000256269 | TRUE | TRUE | TRUE | TRUE | 5 | 778 | TRUE | TRUE | 322 |

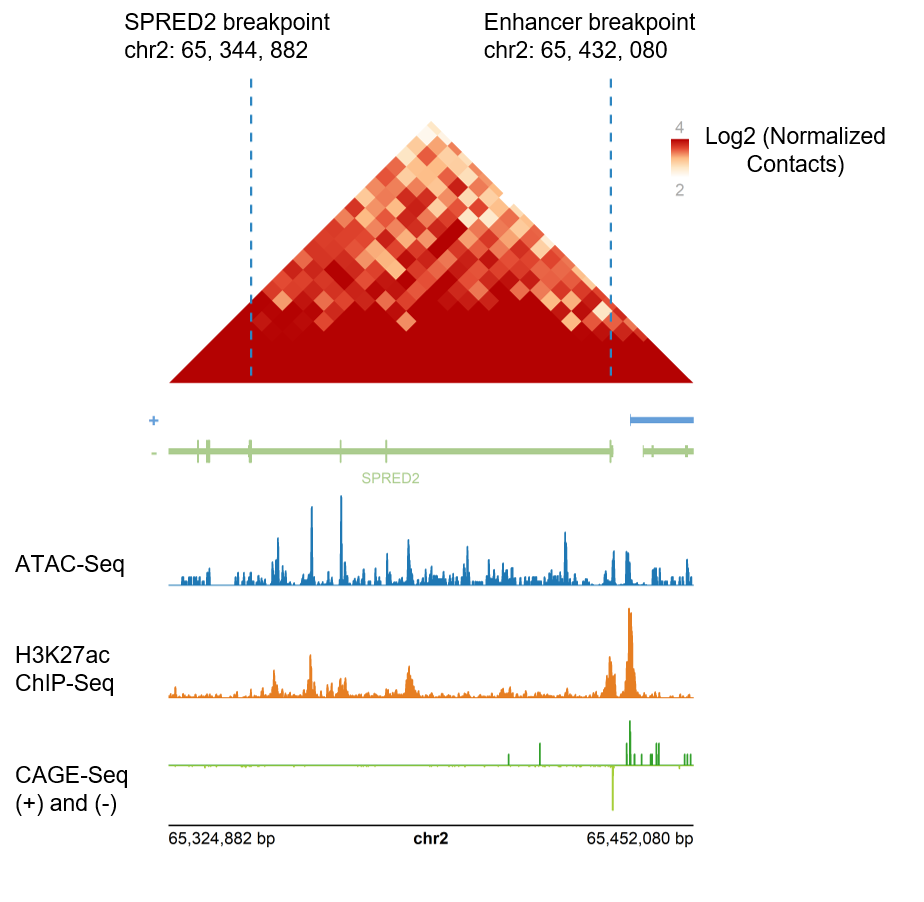

**Supplementary Figure 1. Spatial and transcriptional validation of chimeric eRNA fusion events in K562 cell line.**

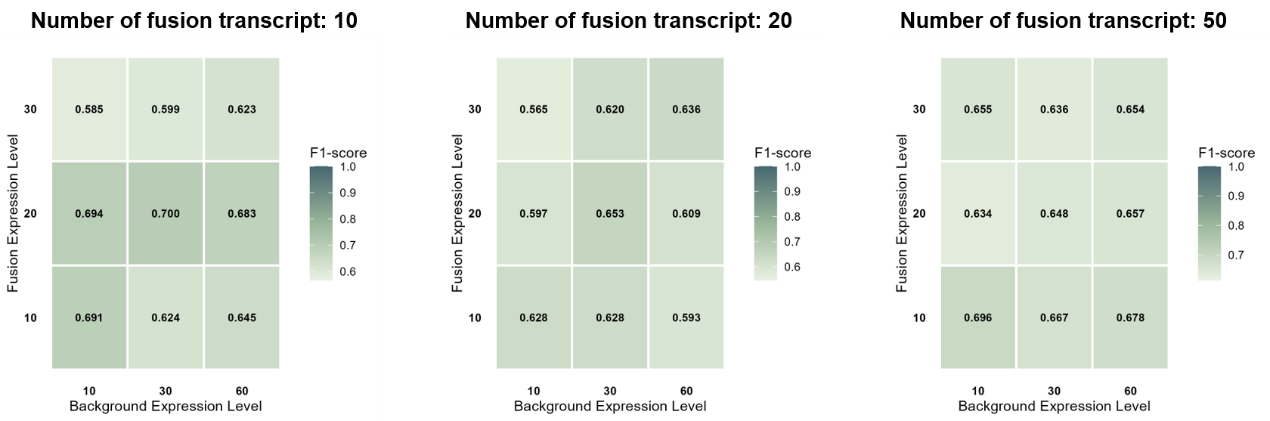

**Supplementary Figure 2. Evaluation of ChiMER robustness and F1-score across diverse parameter configurations using positive control benchmarks.**
